# Computation of transcranial magnetic stimulation electric fields using self-supervised deep learning

**DOI:** 10.1101/2021.11.09.467946

**Authors:** Hongming Li, Zhi-De Deng, Desmond Oathes, Yong Fan

## Abstract

Electric fields (E-fields) induced by transcranial magnetic stimulation (TMS) can be modeled using partial differential equations (PDEs). Using state-of-the-art finite-element methods (FEM), it often takes tens of seconds to solve the PDEs for computing a high-resolution E-field, hampering the wide application of the E-field modeling in practice and research. To improve the E-field modeling’s computational efficiency, we developed a self-supervised deep learning (DL) method to compute precise TMS E-fields. Given a head model and the primary E-field generated by TMS coils, a DL model was built to generate a E-field by minimizing a loss function that measures how well the generated E-field fits the governing PDE. The DL model was trained in a self-supervised manner, which does not require any external supervision. We evaluated the DL model using both a simulated sphere head model and realistic head models of 125 individuals and compared the accuracy and computational speed of the DL model with a state-of-the-art FEM. In realistic head models, the DL model obtained accurate E-fields that were significantly correlated with the FEM solutions. The DL model could obtain precise E-fields within seconds for whole head models at a high spatial resolution, faster than the FEM. The DL model built for the simulated sphere head model also obtained an accurate E-field whose average difference from the analytical E-fields was 0.0054, comparable to the FEM solution. These results demonstrated that the self-supervised DL method could obtain precise E-fields comparable to the FEM solutions with improved computational speed.

## Introduction

Transcranial magnetic stimulation (TMS) is a noninvasive brain stimulation method used in treating major depression and other neuropsychiatric disorders (O’Reardon et al., 2007). However, TMS treatment outcomes vary greatly across patients (Cash et al., 2021; Cash et al., 2020; Diekhoff-Krebs et al., 2017; Fox et al., 2012; Fox et al., 2013; Kim et al., 2014; Luber et al., 2017; Opitz et al., 2016; Sack et al., 2009; Weigand et al., 2018; Williams et al., 2021). One primary source of such variability is that TMS suffers from targeting inaccuracies (Julkunen et al., 2009; Weiss et al., 2013). Recent studies have demonstrated that optimizing TMS stimulation parameters, such as location and orientation of the TMS coil, might improve TMS targeting and focality for individual subjects (Gomez et al., 2021; Makarov et al., 2020b; Weise et al., 2020), and optimizing TMS coil placement based on individual functional neuroanatomy could potentially increase effect sizes for both basic and clinical studies (Diekhoff-Krebs et al., 2017; Fox et al., 2012; Fox et al., 2013; Kim et al., 2014; Luber et al., 2017; Opitz et al., 2016; Weigand et al., 2018). Modeling of electric-fields (E-fields) is now the most widely used method to characterize the localization and spread of electrical current in the brain induced by TMS (Bungert et al., 2017; Deng et al., 2013; Goetz and Deng, 2017; Gomez-Tames et al., 2020; Saturnino et al., 2019; Wang and Eisenberg, 1994). A variety of numerical computational methods have been developed to compute E-fields in conjunction with realistic head models by iteratively solving PDEs governing the E-field induced by TMS coils, including finite element methods (FEMs), boundary element methods (BEMs), and finite-difference methods (FDMs) (Htet et al., 2019; Makarov et al., 2020a; Nielsen et al., 2018; Paffi et al., 2015; Saturnino et al., 2019). The convergence of these numerical methods to the actual solution is sensitive to density of the head mesh, the polynomial approximation order, and error tolerance, and their computational cost is proportional to the modeling accuracy (Babuska et al., 1981). Although compromise is often necessary in real applications, it often takes tens of seconds for state-of-the-art E-field modeling methods to compute a high-resolution E-field (Htet et al., 2019; Makarov et al., 2020a; Nielsen et al., 2018; Paffi et al., 2015; Saturnino et al., 2019).

The high computational cost of E-field modeling also makes it time-consuming and costly to optimize TMS stimulation parameters since a large number of E-fields have to be explored to identify an optimal solution (Gomez et al., 2021; Makarov et al., 2020b; Weise et al., 2020). To benefit clinical practice using these sophisticated tools, the computational cost of E-field modeling has to be reduced substantially without compromising accuracy. Faster E-field modeling is achievable by using a dipole-based magnetic stimulation profile approach that has to compute a magnetic stimulation profile for each individual subject with several hours of CPU time (Daneshzand et al., 2021) or to compute the E-field only on sparse points or the mean of the E-field in a region of interest (ROI) by leveraging the reciprocity principle (Gomez et al., 2021; Koponen et al., 2019). Recent studies have demonstrated that superfast high-resolution E-field modeling can be achieved using deep neural networks (DNNs) (Xu et al., 2021; Yokota et al., 2019). Particularly, the magnitude of E-fields was estimated based on individualized MRI head scans and TMS coil positions by DNNs (Yokota et al., 2019), and 3D vector E-fields were predicted using deep DNNs by integrating both individualized neuroanatomy (scalar-valued tissue conductivity or anisotropic conductivity tensors) and primary E-fields generated by TMS coils as the input (Xu et al., 2021). Though promising results have been obtained, the existing deep learning (DL) based models were driven and optimized in a supervised learning setting. To train the DNNs, E-fields estimated by conventional numerical methods, such as FEM, are used to generate training data. Therefore, their accuracy would be bounded by the conventional numerical methods used to generate the training data. Inspired by self-supervised deep learning methods (Geneva and Zabaras, 2020; Guo et al., 2020; Li et al., 2021; Li and Fan, 2018, 2020; Qin et al., 2019; Raissi et al., 2019; Rao et al., 2021; Tian et al., 2020; Winovich et al., 2019; Yang and Perdikaris, 2019; Zhu et al., 2019) and the pioneer deep learning based E-field computation methods (Xu et al., 2021; Yokota et al., 2019), we develop a novel self-supervised deep learning based TMS E-field modeling method to obtain precise high-resolution E-fields. Specially, given a head model and the primary E-field generated by TMS coil, a DL model is built to generate the electric scalar potential by minimizing a loss function that measures how well the generated electric scalar potential fits the governing PDE, from which the E-field can be derived directly. In contrast to the conventional numerical methods that solve the PDEs iteratively, the DL model is built to learn the solution to the PDE directly. In contrast to the existing supervised DL methods, our DL model is trained in a self-supervised manner by minimizing an energy function that solves the governing PDE as a loss function, which does not require any external supervision. The trained DL model could be applied to new subjects and predict their E-fields by one forward-pass computation. We have validated the proposed DL model using both simulated sphere head model and realistic head models, and experimental results have demonstrated that our method can obtain precise E-fields comparable to solutions obtained by a state-of-the-art FEM implemented in SimNIBS v3.1 with improved computational speed.

## Methods

We develop a self-supervised DL model to compute TMS E-fields by directly learning a mapping from the magnetic vector potential of a TMS coil and a realistic head model to the TMS induced E-field so that high-resolution TMS E-fields will be a good estimate of the solution to the governing PDE of TMS E-fields and be computed by one feedforward computation rapidly.

### TMS E-fields modeling

Given a head model consisting of head tissue compartments with different conductivities, the E-field *E* in the head induced by a TMS coil can be computed by solving a PDE (Goetz and Deng, 2017; Gomez-Tames et al., 2020; Wang and Eisenberg, 1994). Based on quasi-static approximation, the E-field, 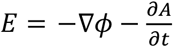, can be computed by solving

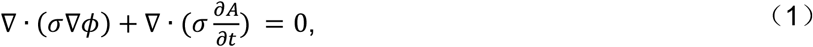

with the Neumann boundary condition

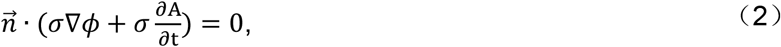

where *σ* is the tissue conductivity, *A* is the magnetic vector potential of the TMS coil, *ϕ* is the electric scalar potential, and 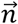 is the normal vector to the tissue surface. Particularly, the primary E-Field 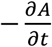 depends only on the TMS coil characteristics (Deng et al., 2013; Koponen et al., 2017) and the secondary field −∇*ϕ* is caused by surface charges in the conducting medium characterized by the head model.

### Computing TMS E-fields using deep neural networks

In contrast to the prevailing FEM/BEM methods adopted in E-field modeling studies (Gomez et al., 2021; Koponen et al., 2019; Makarov et al., 2020a; Makarov et al., 2020b; Nielsen et al., 2018; Paffi et al., 2015; Saturnino et al., 2019; Weise et al., 2020) and the existing DL methods that learn a mapping from individual MRI head scans/anatomy to E-fields in a supervised learning framework (Xu et al., 2021; Yokota et al., 2019), our DL model is built to minimize an energy function that solves the governing equation of Eq. (1) with the boundary condition of Eq. (2) in a self-supervised fashion as illustrated in Fig. 1. Given a head model and TMS induced primary E-field as input, a deep neural network with parameters Θ is built to estimate *ϕ* by minimizing a loss function *L* that measures the dissipated power in the conducting medium (Wang and Eisenberg, 1994), specified as:

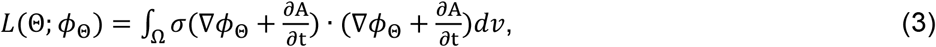

where *ϕ*_Θ_ is the electric scalar potential computed by the deep neural network and *ν* refers to a spatial location (voxel) within the head model Ω.

**Fig.1.**
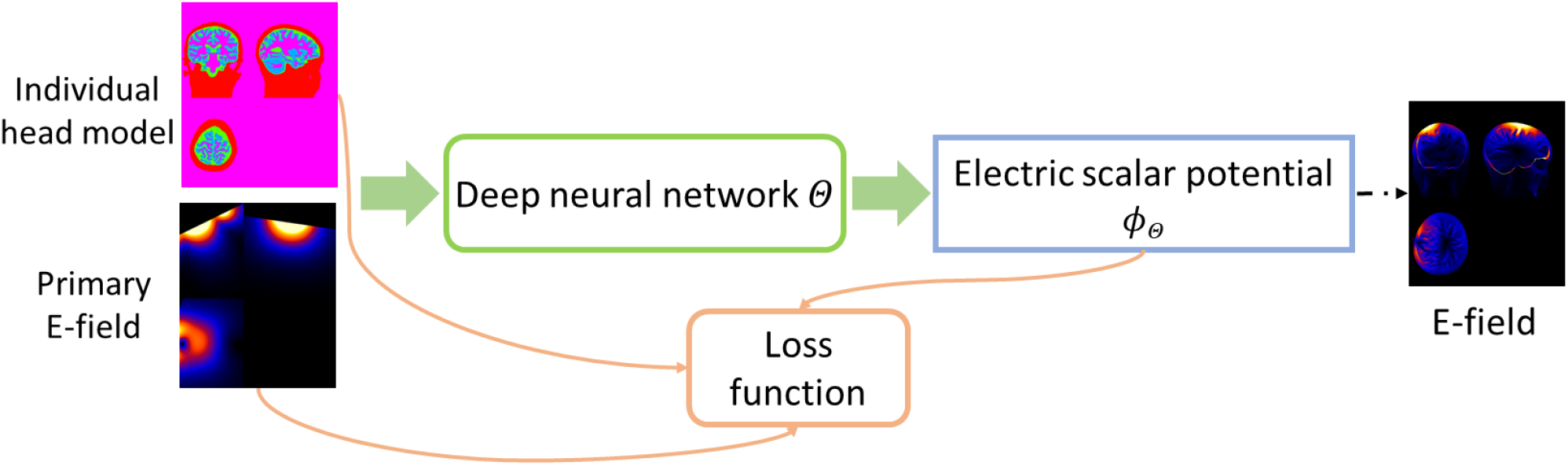
A self-supervised deep learning model for computing TMS E-field. A deep neural network is applied to learn a mapping from individual head model and TMS induced primary E-field to the TMS electric scalar potential, and the network is optimized by a loss function determined by an energy function (Wang and Eisenberg, 1994).

Given the input training data, the deep neural network is trained to optimize the loss function of Eq. (3) in a self-supervised manner. Once the deep neural network is optimized, it could be applied to new subjects and predict the electric scalar potential *ϕ* by one forward-pass computation, from which the E-fields could be computed directly.

### Network architecture for computing TMS E-fields

The overall architecture of our deep neural network for computing TMS E-fields is illustrated in Fig. 2. The deep neural network’s backbone is a U-Net with an Encoder-Decoder architecture (Ronneberger et al., 2015). The network’s input consists of an individual head model (scalar tissue conductivity map, a 4D volume with a channel size of 1) and a subject-specific primary E-field (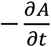, a 4D volume with a channel size of 3), and its output includes the estimated electric scalar potential *ϕ*_Θ_ (a 4D volume with a channel size of 1 and the same spatial dimension as the head model) and its gradient. The total E-field will be estimated as 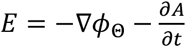. Particularly, the encoder path consists of ten convolutional layers with 8 to 128 filters and a stride of 1 or 2 for downsampling, the decoder path consists of four deconvolutional layers with 128, 64, 32, and 16 filters and a stride of 2 for upsampling, each of which is followed by two additional convolutional layers with 64, 32, 16, and 16 filters and a stride of 1. One output convolutional layer with 1 filter is used to predict the electric scalar potential *ϕ*_Θ_, and its gradient is computed with a central difference operator on the image grid. Leaky ReLU (Maas et al., 2013) activation function is used for all the convolutional and deconvolutional layers, except those two output layers. The kernel size in all layers is set to 3×3×3.

**Fig.2.**
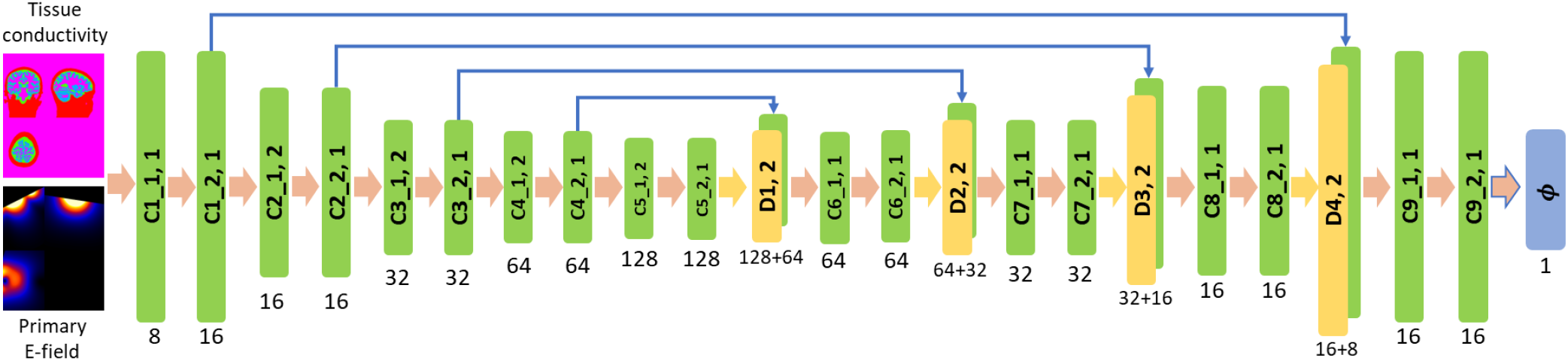
Deep convolutional neural network with an Encoder-Decoder architecture applied to learn TMS E-field. The numbers underneath convolutional (C1_1, C1_2, …, C9_2) and deconvolutional (D1, D2, D3, and D4) layers indicate their corresponding numbers of kernels, with a stride of 1 or 2 for downsampling or upsampling. The kernel size in all layers is set to 3×3×3.

## Experimental results

### Data preparation

We have evaluated the proposed deep learning method using both simulated sphere head model and realistic head models from real MRI scans.

For the simulated data, a 3D sphere head model with a radius of 95 mm and isotropic resolution of 1 mm was generated. Its origin coordinate was set to [0, 0, 0] and its conductivity set to 1 s/m homogeneously. The excitation was given by a point magnetic dipole located outside of the sphere. The dipole’s location was set to [0, 0, 100] and its moment set to [0, 0, 1]. The TMS induced E-field of this sphere head model can be calculated analytically (Heller and van Hulsteyn, 1992), facilitating the direct evaluation of numerical accuracy of methods under comparison.

For the realistic head models, we adopted a local cohort of 125 healthy adult subjects with high-resolution multi-echo T1-weighted MPR images (TR=2400 ms, TI=1060 ms, TE= 2.24 ms, FA=8°, 0.8×0.8×0.8 mm^3^ voxels, image size=208×300×320, FOV= 256 mm). Based on these MRI scans, we used SimNIBS v3.1 to generate anatomically accurate head models (‘headreco’ option with SPM/Computational Anatomy Toolbox for tissue segmentation) and compute primary E-fields induced by a Magstim 70mm Figure-of-Eight coil placed at varied locations with different orientations (Gomez-Tames et al., 2018). For meshes of head models used in FEM computation, the average number of tetrahedrons was 3.915 × 10^6^ (with std of 3.633 × 10^5^), and the average edge length was 2.103 mm (with std of 0.801 mm), as the default setting used in the SIMNIBS pipeline. The tissue conductivity map was generated by substituting the head tissue label values with their corresponding conductivity values. We adopted the SimNIBS conductivity values, i.e., 0.126, 0.275, 1.654, 0.01, 0.465, and 0.5 S/m for whiter matter, gray matter, CSF, bone, scalp, and eyes respectively.

### Experiment settings and implementation

To obtain the E-field of the simulated sphere head model using our proposed deep learning model, the conductivity map of the sphere and the primary E-field induced by the dipole was fed into the network as illustrated in Fig. 2, and the network was trained and optimized with respect to the loss function of Eq. (3) until convergence.

For the evaluation on the realistic head models, we randomly selected 100 subjects as training subjects and the remaining 25 as testing subjects. To evaluate the accuracy and robustness of our deep learning model with respect to different TMS coil positions and directions, three different experiment settings were adopted.

*Setting 1*. The TMS coil was placed at a location within the motor cortex (center=‘C1’ and pos_ydir=‘CP1’ as defined in the EEG10-10 system) for each subject to generate a primary E-field using SimNIBS. There were 100 pairs of subject specific tissue conductivity maps and primary E-fields in total used for training the deep learning model and 25 pairs for testing its accuracy under this setting, which served as a proof-of-concept validation of our deep learning model on realistic head models.

*Setting 2*. The TMS coil was placed at the same location within the motor cortex (center=‘C1’) but in different directions for training and testing the deep learning model. Particularly, primary E-fields of each training subject were generated with different coil directions (seen directions), including ‘CP1’, ‘Cz’, ‘FC1’, and ‘C3’, as training data. There were 400 pairs of subject specific tissue conductivity maps and primary E-fields in total for training the deep learning model. The optimized deep learning model was then evaluated on the testing subjects with the coil placed in directions different from those for generating the training data (unseen directions), including ‘CPz’, ‘FCz’, ‘FC3’, and ‘CP3’. In total, there were 100 pairs of subject specific tissue conductivity maps and primary E-fields for testing. This setting was adopted to evaluate the generalization performance of the deep learning model with respect to varying coil directions.

*Setting 3*. The TMS coil was placed at different spatial locations within the left dorsolateral prefrontal cortex (DLPFC) to evaluate the robustness of the deep learning model with respect to changes of coil locations. Particularly, a target position was defined using the average mean Montreal Neurological Institute (MNI) coordinates (x=−42, y=16, z=28) (Friehs et al., 2020), which was transformed to subject space using SIMNIBS to obtain a subject-specific target position. Then, multiple coil positions and directions were generated within a grid centered at the target position using the SIMNIBS function ‘optimize_tms.get_opt_grid’ with parameters of radius=20, resolution_pos=10, resolution_angle=90, angle_limits=[-180,180], yielding 36 (9 positions by 4 directions) pairs of tissue conductivity map and primary E-field for each subject. In total, there were 3600 pairs of subject specific tissue conductivity maps and primary E-fields for training the deep learning model and 900 pairs of subject specific tissue conductivity maps and primary E-fields for testing its accuracy.

For both the simulated and realistic head models, our deep neural network’s input was a concatenated 4D volume data of the scalar tissue conductivity map (a 4D volume with a channel size of 1) and the primary E-field (a 4D volume with a channel size of 3), and its output included a predicted scalar electric potential (a 4D volume with a channel size of 1) and its gradient. The deep neural network was optimized under each setting respectively on the training data regarding the loss function of Eq. (3). The subject-specific head model and primary E-field was generated using SimNIBS for each subject. For the realistic head models, the input image was cropped (only regions outside the head was cropped) to have a spatial dimension of 208×288×304 to fit the fully convolutional network architecture of our DL model. The channel size was 1 and 3 for scalar image (tissue conductivity map and electric potential) and vector image (primary E-field), respectively.

Our deep learning model was implemented using Tensorflow (Abadi et al., 2016). Adam optimizer (Kingma and Ba, 2014) is adopted to optimize the network, the learning rate was set to 1 × 10^−4^, the batch size was set to 1, and the number of iterations is set to 10000 for the simulated sphere head model, 30000 for realistic head model setting 1, 60000 and 90000 for realistic head model setting 2 and 3 during training. One NVIDIA TITAN RTX GPU with 24G memory was used for training and testing. Training losses of our deep learning models on the simulated sphere head model and realistic head models are shown in Fig. 3, demonstrating the optimized deep learning models reached convergence with the specified parameters.

**Fig.3.**
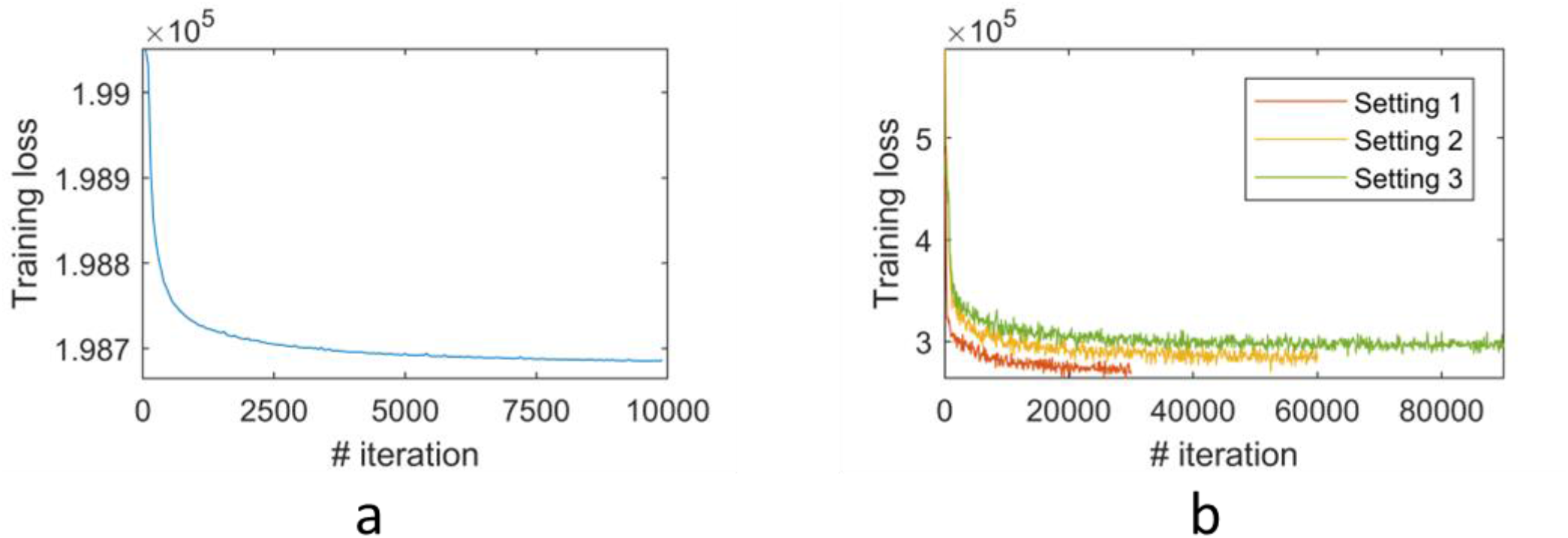
Training loss of the proposed deep learning model on simulated sphere head model (a) and realistic head models under different experiment settings (b).

### Evaluation and comparisons

We compared our method in terms of both accuracy and computational speed with a state-of-the-art FEM, with superconvergent patch recovery, implemented in SimNIBS v3.1. We also compared our method with a FDM under *Setting 1* using an implementation available at https://github.com/luisgo/TMS_Efield_Solvers (Gomez et al., 2020). As no ground truth was available for the realistic head models, we used the solutions of FEM as reference to estimate the accuracy, following the existing deep learning studies of E-field modeling (Xu et al., 2021; Yokota et al., 2019). The FEM solutions were projected onto voxels using “msh2nii” as implemented in SimNIBS for the comparison. Two evaluation metrics, including pointwise magnitude error and correlation coefficient between the predicted and reference solutions were adopted (Gomez et al., 2020). Specifically, the correlation coefficient was computed as Pearson correlation between the magnitude of E-fields obtained by our DL method and the FEM within a specified ROI, and the pointwise magnitude error was computed as 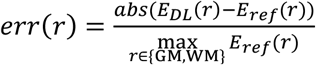, where *r* refers to voxel *r* in GM and WM region. Both measures were evaluated within different ROIs, including the combined gray matter and whiter matter (GM&WM) region, the gray matter (GM) region, the white matter (WM) region, region thresholded at the 95th percentile of E-field magnitude (Xu et al., 2021), and region thresholded at the 50% of the E-field maximum (Deng et al., 2013; Yokota et al., 2019), to get better understanding about the characteristics of the DL based solution. For the simulated sphere head model, the E-field can be calculated analytically (Heller and van Hulsteyn, 1992) to directly evaluate the numerical accuracy of methods under comparison. Particularly, the difference between a numerical solution *E*_*num*_ and an analytically solution *E*_*ana*_ was measured with a normalized root-mean-square error 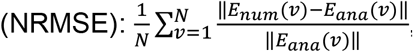, where *N* is the number of voxels in the sphere head model. The evaluation metrics were computed on voxels with values greater than 0 in the reference/ground-truth solution.

## Results

### Results on simulated sphere head model

Fig. 4 shows the magnitude of E-fields computed analytically, by the FEM, and by the proposed DL model, respectively, demonstrating that the numerical E-fields obtained by our DL model and the FEM were visually similar to the analytical solution. The NRMSE between the DL based and analytical E-field was 0.0054, close to the NRMSE between FEM based and analytical E-field (NRMSE=0.0055), indicating that the E-field solution obtained by the DL model was comparable to the FEM solution. This result demonstrated that the proposed DL model can indeed be optimized to learn the E-field that follows the physics law underlying the TMS stimulation.

**Fig.4.**
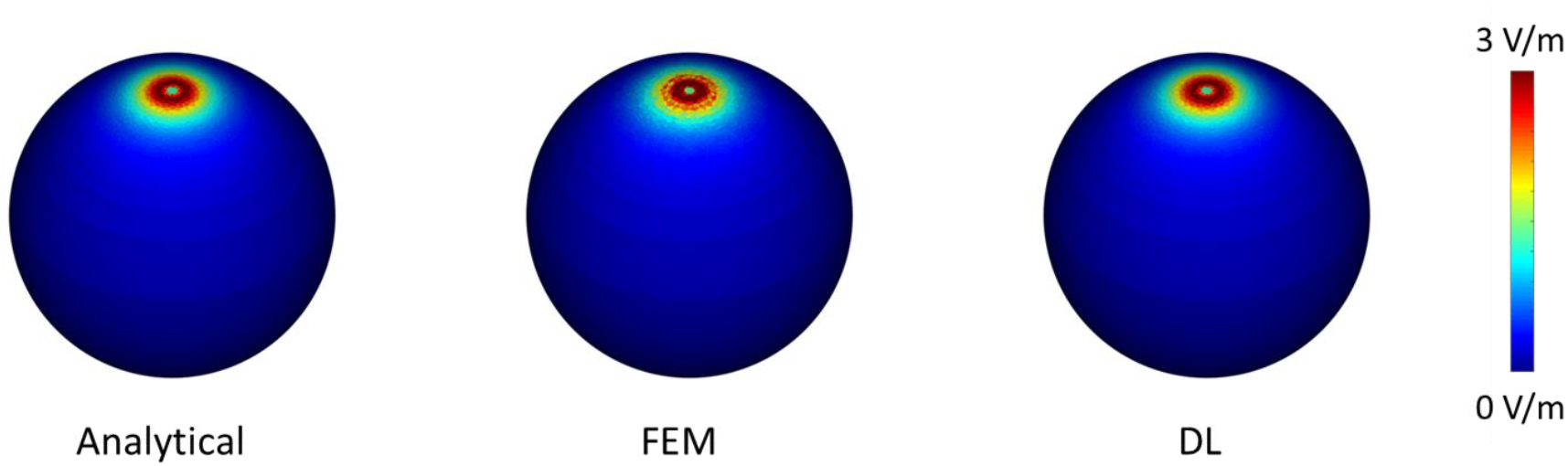
The magnitude of E-fields of the sphere model computed analytically, by FEM (with a NRMSE of 0.0055), and by our DL method (with a NRMSE of 0.0054), respectively.

### Results on realistic head model setting 1

The E-fields of three randomly selected testing subjects computed by the FEM and our proposed DL method are shown in Fig. 5. The results obtained by our DL model had patterns similar to those obtained by FEM. Quantitatively, the results obtained by our DL method were significantly correlated with the FEM solutions, with an average correlation of 0.978 and an average pointwise magnitude error of 0.012 within the gray and white matter region, and these measures were similar when evaluated within gray matter and white matter respectively, as summarized in Table 1, indicating that the results by our DL model are comparable to the reference solutions of FEM. The average correlation and pointwise magnitude error were 0.948 and 0.03 respectively for regions with E-field magnitude exceeding the 95^th^ percentile, which are similar to those obtained by state-of-the-art supervised DL models (Xu et al., 2021). For regions with a E-field magnitude exceeding 50% of the E-field maximum, the average correlation and pointwise magnitude error were 0.837 and 0.049 respectively. These quantitative measures were largely consistent with results shown in Fig. 5 that the magnitude error was relatively larger for regions with a high E-field magnitude. Moreover, our DL model obtained similar correlation coefficients and pointwise magnitude error on both training and testing subjects as shown in Table 1, demonstrating the model’s robustness to the anatomical differences across different subjects. Our E-field solutions were also significantly correlated with the FDM solutions, with an average correlation coefficient of 0.981 (standard deviation: 0.006) and an average pointwise magnitude error of 0.010 (standard deviation: 0.002) within the gray and white matter.

**Fig.5.**
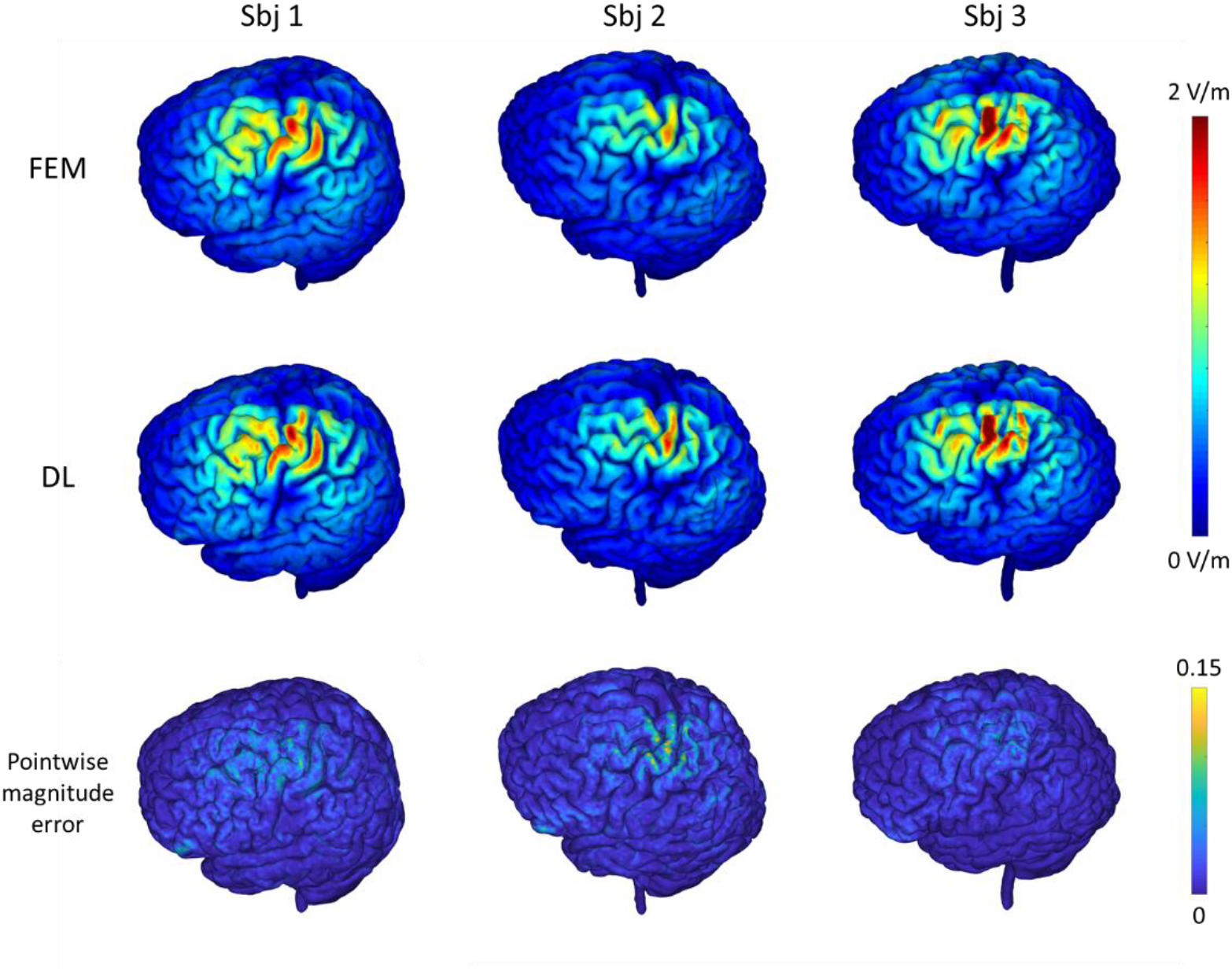
E-fields of three randomly selected testing subjects computed by the FEM and the proposed DL model with the motor cortex as a stimulation target (1st and 2nd rows), and the corresponding pointwise magnitude error map with the FEM solution as reference (3rd row). The top colorbar shows the magnitude values (in V/m) of the E-fields and the bottom colorbar shows the normalized pointwise magnitude error.

**Table 1.** Quantitative evaluation of the proposed DL method under different experimental settings, with the FEM solutions as reference. Mean and standard deviation of the correlation coefficient (CC) and pointwise magnitude error (PME) measures within different regions of interest (ROIs) are demonstrated. ROI definition: gray and whiter matter (GM&WM), gray matter (GM), white matter (WM), region thresholded at the 95th percentile of E-field magnitude (95th percentile), region thresholded at the 50% of maximum E-field magnitude (50% of max).

|  | CC<br>GM&WM | CC<br>GM | CC<br>WM | CC<br>95<sup>th</sup><br>percentile | CC<br>50% of<br>Max | PME<br>GM&WM | PME<br>GM | PME<br>WM | PME<br>95<sup>th</sup><br>percentile | PME<br>50% of<br>Max |
| --- | --- | --- | --- | --- | --- | --- | --- | --- | --- | --- |
| Setting 1 train | 0.984<br>±0.005 | 0.984<br>±0.004 | 0.985<br>±0.006 | 0.958<br>±0.007 | 0.856<br>±0.035 | 0.010<br>±0.002 | 0.010<br>±0.002 | 0.010<br>±0.002 | 0.025<br>±0.003 | 0.044<br>±0.003 |
| Setting 1 test | 0.978<br>±0.006 | 0.979<br>±0.005 | 0.977<br>±0.008 | 0.948<br>±0.014 | 0.837<br>±0.034 | 0.012<br>±0.003 | 0.012<br>±0.003 | 0.013<br>±0.003 | 0.030<br>±0.005 | 0.049<br>±0.008 |
| Setting 2 test (seen<br>dir) | 0.979<br>±0.005 | 0.978<br>±0.005 | 0.980<br>±0.006 | 0.933<br>±0.017 | 0.801<br>±0.127 | 0.013<br>±0.003 | 0.013<br>±0.003 | 0.013<br>±0.003 | 0.034<br>±0.007 | 0.055<br>±0.011 |
| Setting 2 test<br>(unseen dir) | 0.957<br>±0.012 | 0.961<br>±0.010 | 0.953<br>±0.015 | 0.918<br>±0.022 | 0.782<br>±0.113 | 0.019<br>±0.005 | 0.019<br>±0.005 | 0.022<br>±0.005 | 0.042<br>±0.011 | 0.064<br>±0.014 |
| Setting 3 test | 0.984<br>±0.004 | 0.983<br>±0.004 | 0.985<br>±0.004 | 0.943<br>±0.010 | 0.819<br>±0.052 | 0.012<br>±0.002 | 0.012<br>±0.002 | 0.012<br>±0.003 | 0.036<br>±0.005 | 0.055<br>±0.006 |

### Results on realistic head model setting 2

The E-fields of three randomly selected testing subjects with unseen coil directions computed by the FEM and our proposed DL method are shown in Fig. 6. Though the coil directions were different from those used for training the DL model, the DL model still successfully predicted the E-fields for the testing subjects, demonstrating good consistency with those obtained by the FEM. Quantitatively, the results on both seen and unseen coil directions obtained by our DL method were significantly correlated with the FEM solutions, with an average correlation of 0.979 and 0.957 respectively, while the average pointwise magnitude error was 0.013 and 0.019 respectively for the gray and white matter region, as summarized in Table 1. As shown in Fig. 6, the prediction error was relatively larger for regions with a high E-field magnitude. It is worth noting that all these results were obtained for testing subjects which were not used for the model training. These results demonstrated that the DL model was robust under different coil direction settings and could generalize to unseen coil directions. We also observed that the pointwise magnitude error of the 3rd subject was larger compared with others in Fig. 6, which might indicate the model’s robustness to inter-subject anatomical difference compromises moderately when the coil directions were not seen during the model training procedure.

**Fig.6.**
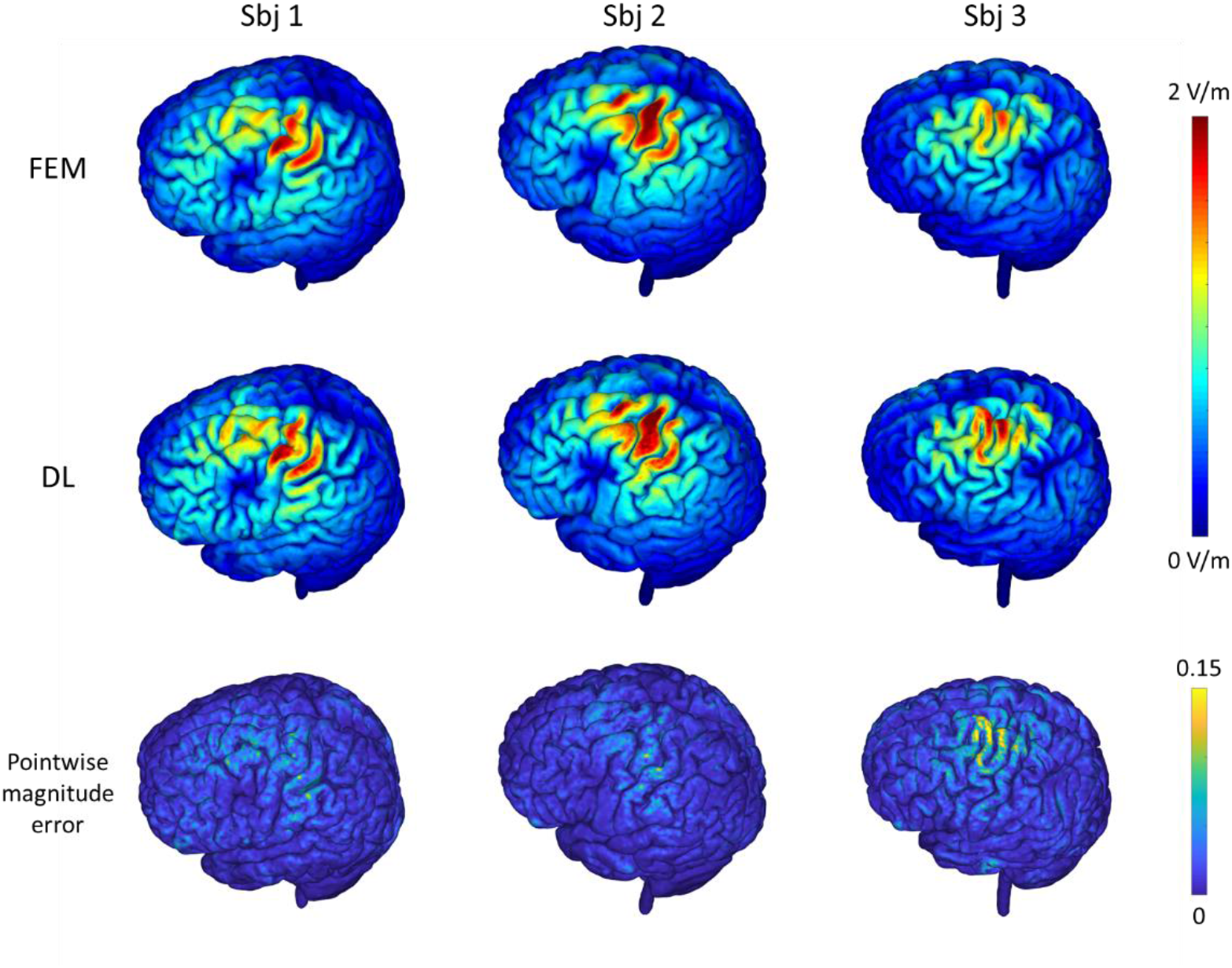
E-fields of three randomly selected testing subjects computed by the FEM and the proposed DL model with the motor cortex as a stimulation target and the coil set in varying directions (1st and 2nd rows), the corresponding pointwise magnitude error map with the FEM solution as reference (3rd row). The top colorbar shows the magnitude values of the E-fields and the bottom colorbar shows the normalized pointwise magnitude error.

### Results on realistic head model setting 3

The E-fields of one randomly selected testing subject computed by the FEM and our proposed DL method for three different coil positions located within the DLPFC area are shown in Fig. 7. The E-fields obtained by our DL model are visually similar to those obtained by FEM. The average correlation coefficient of the solutions obtained by our DL model and FEM was 0.984 within the gray and white matter for the testing subjects, with an average pointwise magnitude error of 0.012 as summarized in Table 1, indicating that our DL model was capable of predicting E-fields with varying coil positions and directions. It could be observed that the prediction error was relatively larger for regions with a high E-field magnitude. The average correlation and pointwise magnitude error were 0.943 and 0.036 respectively for regions with a E-field magnitude exceeding the 95th percentile, and the average correlation and pointwise magnitude error was 0.819 and 0.055 respectively for regions with E-field magnitude exceeding 50% of the E-field maximum.

**Fig.7.**
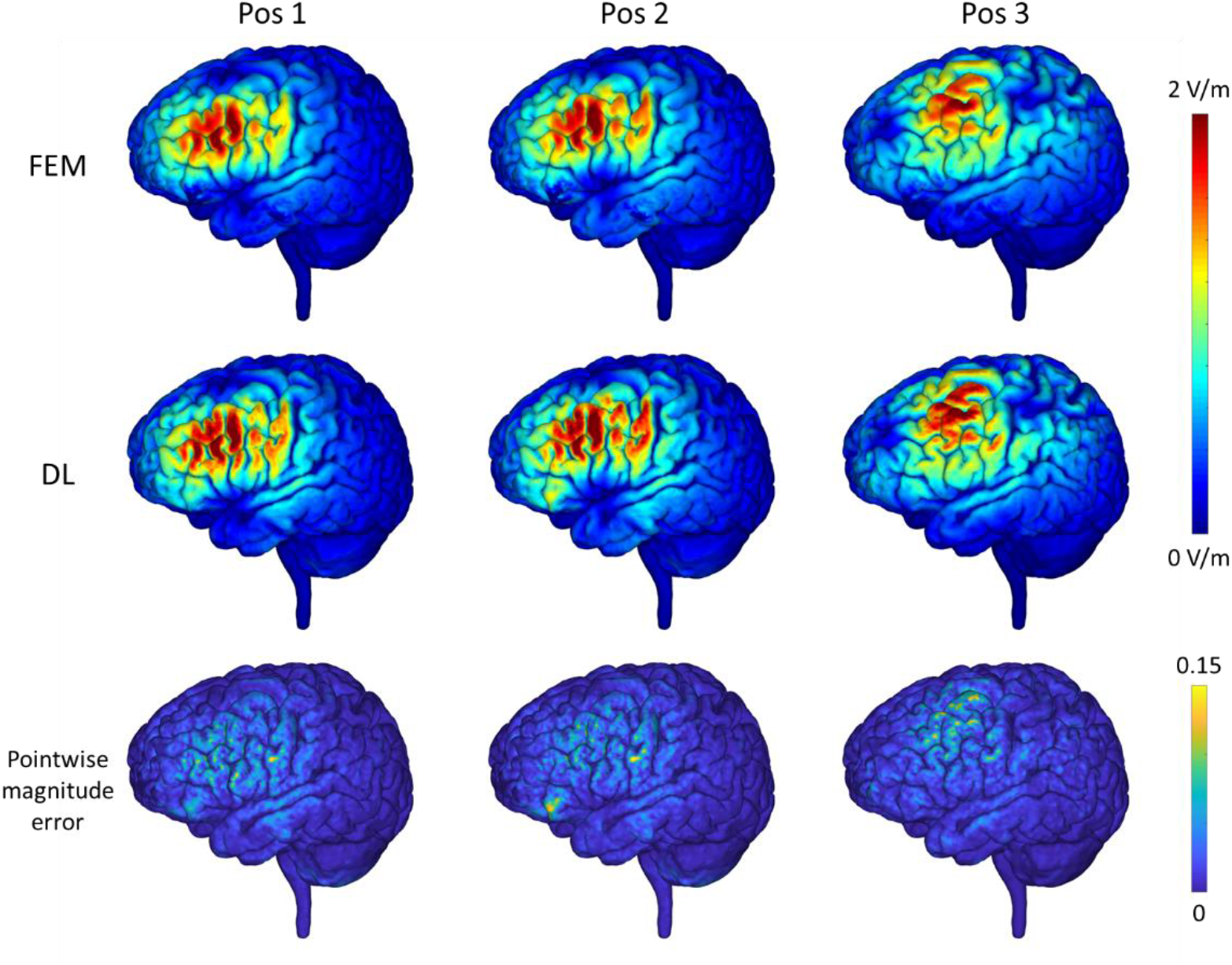
E-fields of one randomly selected testing subjects computed by the FEM and the proposed DL model with the dorsolateral prefrontal cortex (DLPFC) as a stimulation target and the coil set at varying positions and in different directions (1st and 2nd rows), and the corresponding pointwise magnitude error map with the FEM solution as reference (3rd row). The top colorbar shows the magnitude values of the E-fields and the bottom colorbar shows the normalized pointwise magnitude error.

### Computation time

It took 35.17 and 33.65 seconds on average by the FEM as implemented in SIMNIBS to obtain the E-field for one subject (whole head model with 208×288×304 voxels at a spatial resolution of 0.8×0.8×0.8 mm^3^) with the TMS coil located at the motor cortex and DLPFC respectively when using one Intel Xeon Gold 5218 CPU. It took 14.27 and 14.26 seconds by our trained DL model using the same CPU. On one NVIDIA TITAN RTX GPU, it took 1.47 and 1.49 seconds respectively by our trained DL model to compute E-fields with the TMS coil located at the motor cortex and DLPFC, respectively. For the FEM, the timing measurement included the time for assembling and solving the FEM system; For DL method, the timing measurement includes the time for one forward pass to obtain the E-field and the electric potential using a head model and a primary E-field as input. We did not include the time for computing head model/mesh and primary E-field for both the FEM and our DL method. This comparison demonstrated the improved computational speed obtained by the proposed deep learning model. For the model training, it took 2.88 seconds on average for each training iteration. It is worth noting that the model only needs to be trained once for a target ROI, which can be applied to new testing subjects without further optimization once trained.

## Discussion

We have developed a self-supervised deep learning model to directly learn a mapping from the magnetic vector potential of a TMS coil and a realistic head model to the TMS induced E-fields, instead of iteratively solving equations governing the E-field induced by a TMS coil. Experimental results on both simulated and realistic head models have demonstrated that our method could obtain similar accuracy compared with the most commonly used numerical method. Our method is capable of computing the TMS induced E-fields by one forward-pass computation, taking less than 1.5 seconds for a realistic head model with high spatial resolution.

Several numerical computational methods have been developed for accurate E-fields modeling in conjunction with realistic head models, such as FEMs, BEMs, and FDMs. Though the computational speed of state-of-the-art numerical methods has been improved a lot, their computational cost for high-resolution E-fields is still high due to the nature of iterative optimization in their PDE solvers, which compromises their use in the optimization of TMS stimulation parameters in both basic and clinical studies. Instead of solving the governing PDEs from scratch, recently studies have demonstrated promising performance of deep neural networks for rapid estimation of E-fields (Xu et al., 2021; Yokota et al., 2019), in which deep neural networks are trained to directly predict the E-fields with high fidelity to those estimated using conventional E-field modeling methods, such as FEMs. Therefore, the deep neural networks are actually trained to predict the solutions obtained by the conventional E-field modeling methods and their performance is bounded by the conventional E-field modeling methods used for generating training data. Moreover, it will also be time-consuming to generate surrogate training data with different TMS stimulation parameters on a large cohort.

In contrast to the existing deep learning based E-field modeling methods (Xu et al., 2021; Yokota et al., 2019) that learn a mapping from head scans/models to surrogate E-fields estimated using conventional E-field modeling methods in the supervised learning way, our method directly learns a solution to the governing equations in a self-supervised learning way, which does not require any external supervision. Our proposed deep neural network is designed to predict the TMS induced electric scalar potential and is optimized so that the network’s output fit the governing PDE as much as possible, which is formulated to minimize an energy function that solves the governing PDE. Therefore, our method directly learns a solution to the same governing PDE as the convention numerical optimization methods do, while benefiting from the fast inference of deep neural networks. As surrogate E-fields are not required by our method, realistic head models from diverse imaging datasets can be used as the training data for our method. To the best of our knowledge, our method is the first study to investigate self-supervised deep learning for TMS E-field modeling, though physics-informed deep learning methods have been successfully applied to solving PDEs on varied domains (Geneva and Zabaras, 2020; Guo et al., 2020; Qin et al., 2019; Raissi et al., 2019; Rao et al., 2021; Tian et al., 2020; Winovich et al., 2019; Yang and Perdikaris, 2019; Zhu et al., 2019).

Our method is designed as a general-purpose method for the computation of E-field, and it could obtain the E-field for a new subject (whole head model with 208×288×304 voxels at a spatial resolution of 0.8×0.8×0.8 mm^3^) in about 1.47 seconds. Several alternative fast E-field computation methods have been developed (Daneshzand et al., 2021; Stenroos and Koponen, 2019). Particularly, a magnetic stimulation profile for each subject needs to be computed in advance (Daneshzand et al., 2021), which may require hours of CPU time. The surface integral equation used in (Stenroos and Koponen, 2019) is valid only for isotropic medium, which cannot account for the anisotropic conductivity of white matter properly. Our method does not require per-subject training or pre-computation, it could be applied to new subjects directly without further optimization once a DL model training is finished. Moreover, we can further reduce its computation time by applying the trained model to a region of interest (ROI) instead of the whole head model, facilitated by the fully convolutional network architecture of our method. It should be noted that an isotropic tissue conductivity model was adopted in our current method development and evaluation, and the trained model should not be applied to head models with anisotropic tissue conductivity. It should be also noted that extension to anisotropic tissue conductivity is feasible by incorporating anisotropic conductivity tensor into the loss function defined in Eq. (3) and replacing the scalar conductivity with anisotropic conductivity tensor properly as the network input.

In addition to promising accuracy and computational speed, our method is robust to varying TMS coil locations and directions. As demonstrated in Fig. 6, the deep neural network generalized well for computing E-fields of the testing subjects that were generated by the coil placed in directions different from those for generating the training data. The validation experiment with the coil placed at left DLPFC has further demonstrated that the accuracy of the predicted E-fields was still comparable to that obtained by the FEM on the testing subjects, even though the coil was placed at varying locations and in different directions, as illustrated in Fig. 7. The good generalization performance of the proposed method may be attributed to its self-supervised learning nature, which optimizes the deep learning model to learn the underlying mapping between head anatomy and electric potential without external guidance or prior assumptions. The robust generalization emphasizes that this method is likely to be applicable to the optimization of TMS stimulation where varying positions and directions around the target position are to be explored. Nevertheless, it was observed in Figs. 5 to 7 that the pointwise magnitude errors were relatively large at the regions with a high E-field magnitude, indicating there is still room for improving the model’s performance with respect to both methodological development and model training, such as optimizing the neural network architecture and increasing the size of training dataset. It was also observed in Fig. 6 that the prediction error for the 3rd subject was relatively larger than that for other subjects, which might be due to large inter-subject anatomical differences between training and testing subjects, and it is expected that a model trained on a larger training dataset with more subjects will improve the model’s robustness to anatomical differences across subjects. On the other hand, the DL-based results can also be served as a good initialization for conventional PDE solvers to achieve an improved convergence rate given that the DL-based results were close to the FEM-based solutions.

Though the self-supervised deep learning method has demonstrated enormous potential for fast and accurate E-field modeling, there are still several limitations should be noted. First, current evaluation was focused on E-field induced by a Figure-of-Eight coil and head models at a single resolution, the influence of different coils and spatial resolutions of head models merits further investigation. Second, scalar tissue conductivity maps were used in the present study. Future work will be devoted to modeling of anisotropic tissue conductivity with deep learning and exploring its effects on the derived E-field. Third, our current model adopts a traditional U-Net architecture, which can be optimized in terms of both accuracy and computational speed of the E-field modeling using neural architecture search (NAS) techniques (Elsken et al., 2019). In addition, tuning the hyper-parameters for network training such as batch size along with the architecture optimization may further improve the performance. Moreover, the gradient operator currently used to compute the loss function in Eq. (3) was implemented as a central difference operator on the image grid, which may generate blurring or artificially elevated peak values around the tissue boundaries. More attention should also be paid to exploring other numerical solutions such as cubic spline based method and spectral differentiation for computing gradient or other processing strategies to improve the prediction at tissue boundary regions. Fourth, dedicated deep neural networks were trained separately for different target regions in the present study, future work will be devoted to investigating the feasibility of one unified neural network for multiple target regions across the cerebral cortex and its generalization with respect to the size of training data and the neural network capacity.

In conclusion, a self-supervised deep learning model was developed to estimate TMS induced E-fields directly from realistic head models and the TMS coil’s magnetic vector potential. The DL model can obtain high-resolution E-fields from realistic head models with high accuracy, facilitating fast and precise TMS E-field modeling, and therefore the optimization of TMS stimulation parameters in both basic and clinical studies.

## Acknowledgements

We sincerely thank the reviewers for their insightful comments and constructive suggestions, which helped improve the manuscript substantially. This work was supported in part by the National Institutes of Health [grant numbers: MH120811, EB022573, and AG066650].

